# The evolutionary history of the current global *Ramularia collo-cygni* epidemic

**DOI:** 10.1101/215418

**Authors:** Remco Stam, Hind Sghyer, Martin Münsterkötter, Saurabh Pophaly, Aurélien Tellier, Ulrich Güldener, Ralph Hückelhoven, Michael Hess

## Abstract

Ramularia Leaf Spot (RLS) has emerged as a threat for barley production in many regions of the world. Late appearance of unspecific symptoms caused that *Ramularia collo-cygni* could only by molecular diagnostics be detected as the causal agent of RLS. Although recent research has shed more light on the biology and genomics of the pathogen, the cause of the recent global spread remains unclear.

To address urgent questions, especially on the emergence to a major disease, life-cycle, transmission, and quick adaptation to control measures, we de-novo sequenced the genome of *R. collo-cygni* (urug2 isolate). Additionally, we sequenced fungal RNA from 6 different conditions, which allowed for an improved genome annotation. This resulted in a high quality draft assembly of about 32 Mb, with only 78 scaffolds with an N50 of 2.1 Mb. The overall annotation enabled the prediction of 12.346 high confidence genes. Genomic comparison revealed that *R. collo-cygni* has significantly diverged from related *Dothidiomycetes*, including gain and loss of putative effectors, however without obtaining species-specific genome features.

To evaluate the species-wide genetic diversity, we sequenced the genomes of 19 *R. collo-cygni* isolates from multiple geographic locations and diverse hosts and mapped sequences to our reference genome. Admixture analyses show that *R. collo-cygni* is world-wide genetically uniform and that samples do not show a strong clustering on either geographical location or host species. To date, the teleomorph of *R. collo-cygni* has not been observed. Analysis of linkage disequilibrium shows that in the world-wide sample set there are clear signals of recombination and thus sexual reproduction, however these signals largely disappear when excluding three outliers samples, suggesting that the main global expansion of *R. collo-cygni* comes from mixed or clonally propagating populations. We further analysed the historic population size (Ne) of *R. collo-cygni* using Bayesian simulations.

We discuss how our genomic data and population genetics analysis can help understand the current *R. collo-cygni* epidemic and provide different hypothesis that are supported by our data. We specifically highlight how recombination, clonal spreading and lack of host-specificity could further support global epidemics of this increasingly recognized plant disease and suggest specific approaches to combat this pathogen.

## Introduction

Plant pathogens cause serious damage on crop plants. They need to be controlled to achieve sufficient yield and best quality of the agricultural products. The ascomycete fungus *Ramularia collo-cygni* has been detected in barley samples worldwide [1]. The filamentous fungus is the major biotic agent of a leaf spotting complex [2] typically occurring late in the growing season on the upper canopy [3]. Ramularia leaf spot (RLS) poses a major risk in barley production, particularly in important barley growing regions like Scotland, mid Europe, Argentina and Uruguay It is estimated to cause losses up to 25 and in extreme cases up to 70% of the yield potential through a significant decrease of kernel size and quality [4]. Since thus far no major resistance genes were identified within the commercial barley genepool, control has been relying on the pre-epidemic use of several fungicides, but only a limited number of active substances are available and resistance has already been reported [5]. *Ramularia collo-cygni* was first described in 1893 by Cavara [6] as *Ophiocladium hordei*. However, it is only since the mid-1980’s that it became of increasing importance, with serious economic impact and the reasons for this remain unknown. Little is know about the pathogens biology or diversity in the field and even reproductive mode in the field and methods of dispersal remain largely unknown.

Previously a draft genome had been published for *R. collo-cygni* strain DK05 [7]. The assembled 30.3 Mb genome data predicted 11,617 of gene models. It allowed the confirmation that *R. collo-cygni* belongs to the *Dothidiomycetes*, particularly to the family of *Mycosphaerellaceae* that contains several major plant pathogens. Additionally the study revealed relative paucity of plant cell wall degrading enzyme genes and a large number of secondary metabolite production associated genes. Both findings were hypothesised to be linked to the relatively long asymptomatic growth inside the host. The authors also highlighted the occurrence of several toxin encoding genes, on the genome, including the well-studied rubellins[8,9]. It is suggested that RLS-associated necrosis is caused by toxins produced by the fungus[9,10]. However, these toxins are non-host-specific and also present in other *Dothideomycetes*, thus their findings did not explain the recent emergence of the pathogen in barley.Instead it has been suggested that the intensive use of mildew resistant *mlo*-genotype cultivars might be one of the causes of the emergence of *R. collo-cygni* as a threat to barley production and quality. McGrann *et al* have investigated the trade-off between the barley *mlo* mutation-mediated powdery mildew resistance and susceptibility to RLS [11]. However, analysis of grain of near isogenic *MLO* and *mlo*-barley did subsequently not suggest enhanced levels of Rcc transmission via *mlo*-barley seed [12].

*R. collo-cygni* has also been isolated from wheat, oat, maize and from a number of uncultivated grasses such as *Agropyron repens* suggesting a broad host range [3] and shows microscopically similar compatible interactions with many of these grasses [13]. It is so far not known whether isolates of *R. collo-cygni* generally are able to infect a broad range of host species or if host specialization (e.g. to barley) occurs within local populations, nor do we have good insights in geographical diversity. An Amplified Fragment Length Polymorphism study investigating the population structure of samples from barley in Czechia, Slovakia, Germany and Switzerland showed variability between the samples and rejected the hypothesis of random mating in the field, but suggests that mixed reproduction is likely [14]. To date, no teleomorph of *R. collo-cygni* has been described. However data from both Amplified Fragment Length Polymorphism or microsatellites studies [15,16] also hint that sexual reproduction might take place in the field. Moreover, both studies show considerable variation between their two samples locations (Scotland & Denmark or Scotland & Czechia, respectively), but do not see clear substructures within the populations.

Here we move beyond the genomics of *R. collo-cygni* and combine a comparative genomics approach with population genetics to get a deeper insight in *R. collo-cygni’*s biology and to better understand the life cycle as basis for sustainable control. We generated an independent draft genome and use this reference to gain a deeper understanding of the pathogens genome compared to selected model fungi and economically relevant plant pathogens, to identify those genetic factors that might contribute to its recent success. Moreover, we sequenced 17 additional isolates from a global collection from different hosts, to infer the pathogen’s population structure and its global diversity. Finally, we used our high quality assembly to infer the Linkage Disequilibrium, to define whether indeed *R. collo-cygni* is a sexual plant pathogen and to establish the historic population sizes of the pathogen. Our combined results show that *R. Collo-cygni* is unlike many other pathogens and that special care might be required to prevent worsening of the current epidemic.

## Materials and Methods

### Fungal isolates and culture maintenance

The isolate Urug2 (isolated in our laboratory from barley leaves collected in Uruguay) was used for the genome sequencing and expression analysis. It was stored in screw-cap test tubes that contained sterilized one fourth strength potato dextrose broth (¼PDB) (Carl Roth GmbH + Co. KG). Mycelium of the isolate was transferred to one fourth strength potato dextrose agar (¼PDA) (Carl Roth GmbH + Co. KG) that was contained in 90-mm-diameter plastic petri dishes.

### *R. collo-cygni Archive samples and* DNA quantification

Archive barley seed samples dating back to the 60’s were provided by Markus Herz from the Bavarian State Research Center for Agriculture (LfL). Quantification of *R. collo-cygni* DNA from archive barley grains sample was performed following a Taq-Man qPCR protocol. Briefly, 1x iQ Supermix (Bio-Rad, Hercules, CA), 400 nmol^−1^ forward and reverse primer (RamF6/RamR6), 150 nmol^−1^ Ramularia probe (FAM; Ram6), 5 μl of DNA template (20 ng μl^−1^), and PCR-grade water were mixed together to make the reaction mixture with a final volume of 25 µl. PCR amplification was performed using a MX3000P Cycler (Stratagene).

### Cultures media and incubation conditions

*Ramularia collo-cygni* cultures were maintained 9-cm petri dishes incubated at room temperature in the dark.For the genome sequencing, the isolate was grown on Alkyl Esther medium (AE) (10 g Yeast extract, 0.5 g MgSO4 *7 H2O, 6 g NaNO3, 0.5 g Kcl, 1.5 g KH2PO4, 20 g Bacto Agar, 20 mL Glycerol for 1000 ml of final volume) for 7 days.

Concerning the expression analysis (by RNA-seq), the isolate was grown on 6 different conditions: “control” which is the same as used for the genome sequencing, *i.e* growing on AE medium for 7 days; “old”, growing the isolates on AE for 14 days instead of 7 days.; “PH5”, growing it on AE buffered with HCl until pH = 5, for 7 days; “PH9”, growing it on AE buffered with NaOH until pH = 9, for 7 days; “NoG”, growing it on AE without glycerol, for 7 days; “BSagar”, growing it on Barley Straw Agar medium (BSagar) (40g of grounded barley straw, 20 g of agar for 1000 mL of final volume), for 7 days.

### DNA and RNA extraction

Mycelia from above-mentioned cultures were harvested, and finely ground mycelium was used for DNA extraction and RNA extraction.

For the DNA extraction, 1400µl of preheated (60°C) CTAB extraction buffer (100 mM Tris–HCl, pH 8.0; 20 mM EDTA, pH 8.0; 1.4M NaCl; 2% CTAB; 0.2% β-mercaptoethanol) was added to a 2 ml Eppendorf tube. The tubes were mixed thoroughly and incubated at 60°C for 1h. The tubes were centrifuged for 5 min at 13,000 rpm. One thousand microliter of the supernatant was transferred into a fresh 2 ml tube and then 1 volume of phenol/chloroform/isoamylalcohol (25:24:1) was added and the tubes were gently mixed for 5 to 10 min. Tubes were centrifuged for 20 min at 13,000 rpm and the supernatant was transferred to a new tube. RNase (20µg/ml, DNase-free) was added and the tubes were incubated at 37°C for 30 min. One volume of phenol/chloroform/isoamylalcohol (25:24:1) was added and the tube were gently mixed for 5 to 10 min. Tubes were centrifuged for 20 min at 13,000 rpm and the supernatant was transferred to a new tube. DNA was precipitated by adding 0.1 volume of sodium acetate (3M, pH 5.2) and 0.6 volume of cold isopropanol. The tubes were kept at −20°C overnight. Tubes were then centrifuged for 10 min at 13,000 rpm at 4°C. The supernatant was discarded and the pellet was washed with 500 µl cold 70% ethanol, the pellet was spin down at 13,000 rpm for 1 min at 4°C and the supernatant was discarded. This last step with the ethanol was repeated one more time. The precipitate was dried for 20–30 min at 37°C and then resuspended in 100 μl of sterile Tris-Buffer (10 mM Tris–HCl, pH 8.5).

Total RNA was extracted from *R. collo-cygni* mycelium from the 6 different conditions using TRIzol reagent (Invitrogen, Karlsruhe, Germany). RNA integrity was confirmed using a 2100 Bioanalyzer RNA Nanochip with the RNA 6000 Nano Kit (5067-1511) following the manufacturer’s protocol (Agilent, Santa Clara, CA, USA).

For the isolation of genomic DNA from barley grain, a subsample of 50 g was ground to fine powder using a laboratory mill. DNA extraction from ground Barley seeds was performed following recommendations of the European Community Reference Laboratories (Joint Research Centre 2007) for the isolation of maize DNA, with some modifications described in [22].

### Library preparation and sequencing

Whole genome sequencing of *R. collo-cygni* was performed by Eurofins Genomics GmbH, Ebersberg, Germany, using a short distance library (SD) by fragmentation and end repair of DNA (customized insert size of approx. 500 bp) on the Illumina MiSeq v3 (paired-end sequencing 2=150 bp) and a long jumping distance library (LJD) with a jumping distance of 8 kbp on the Illumina HiSeq 2000/2500 v3 (paired-end sequencing 2=300 bp).

The RNA-seq library preparation was performed using the TruSeq Stranded mRNA Library Prep Kit (RS-122-2101) (Illumina San Diego, California 92122 U.S.A.) following the manufacturer’s protocol. The prepared library (Insert size: about 180 bp) was then subjected to high-throughput sequencing using the Illumina HiSeq2500 using 1 lane and multiplexed paired-end read (read 1: 101 cycles, index read: 7 cycles, read 2: 101 cycles). The sequencing was performed using the TruSeq Rapid PE Cluster Kit (PE-402-4001) and the TruSeq Rapid SBS Kits - HS (200 cycles) (FC-402-4001) (Illumina San Diego, California 92122 U.S.A.).

### Genome assembly and structural annotation

The assembly was performed by ALLPATHS-LG [17] using around 100 fold SD and 30 fold LJD genome covering. The assembly in total has a size of 32 Mb and consists of 78 scaffold with a N50 of 2.1 Mb. Gene models were generated by three de-novo prediction programs: 1) Fgenesh [18] with different matrices (trained on *Aspergillus nidulans*, *Neurospora crassa* and a mixed matrix based on different species); 2) GeneMark-ES [19] and 3) AUGUSTUS [20] with RNA-seq based transcripts as training sets. Annotation was aided by exonerate [21] hits of protein sequences from *Botrytis*, *Sclerotinia* and *Rhynchosporium* models to uncover gene annotation gaps. Transcripts were assembled on the RNA-seq data sets using Trinity[22]. The different gene structures and evidences (exonerate mapped transcripts and bowtie mapped RNA-seq reads) were visualized in Gbrowse [23], allowing manual validation of coding sequences. The best fitting model per locus was selected manually and gene structures were adjusted by splitting or fusion of gene models or redefining exon-intron boundaries if necessary. tRNAs where predicted using tRNAscan-SE [24]. The predicted protein sets were searched for highly conserved single (low) copy genes to assess the completeness of the genomic sequences and gene predictions. All of the 248 core-genes commonly present in higher eukaryotes (CEGs) could be identified by Blastp comparisons (e-value: 10^−3^) [25].

### RNA-seq analysis

RNA-seq reads were mapped against our reference genome using TopHat (ver. 2.1.0). The gene expression levels were estimated as Fragments Per Kilobase of Million Fragments Mapped (FPKM) using the cufflinks.cuffdiff package (ver. 2.2.1). All the FPKM values were transferred into log2 values after adding 1 to each FPKM. Fold changes of expression levels between the different conditions were then calculated. The transcripts with a fold change of expression between two conditions greater or equal to 4 (log2>=2) were considered significantly differentially expressed

### Functional annotation of predicted open reading frames and data repositories

The protein coding genes were analyzed and functionally annotated using the PEDANT system[26] genome and annotation data was submitted to the European Nucleotide Archive (ENA, <u>http://www.ebi.ac.uk/ena/data/view/FJUY01000001-FJUY01000078</u>).

### Prediction of secreted proteins and effectors

We used a pipeline which combines the use of five methods. First, secreted proteins candidates are selected by SecretomeP [27] (cutoff score = 0.6). Proteins with mitochondrial target were removed using TargetP (RC-score < 4) [28] and the one with membrane bound using TMHMM [29]. Extracellular target compartments were predicted using WolfPsort [30]. Finally, to identify the class of secretion (classical or non-classical secreted proteins) SignalP (s-score ≥ 0.5) [31] was applied. The combination of all these 5 methods generated our working set of secreted proteins (Table S2)

For the prediction of effectors, the subsequent set of secreted proteins predicted as explained above was submitted to EffectorP 1.0 [32] (Table S2).

### Species phylogeny and divergence time calculations

Orthologous genes for 11 main House Keeping Genes we selected from the proteomes of all 12 species available from the FunyBASE [33]. Using the Mega7 software suite [34] a ClustalW pairwise Alignmend and multiple aligment with standard parameters for the 10 orthologs combined proteins of the other species (An, Bc, Bg, Cg, Ds, Ff, Fg, Lb=out, Mf, Mo, Pi=out, Pr, Pt, Rc, Sn, Um= outgroup, Zt) was performed.

The evolutionary history was inferred by using the Maximum Likelihood method based on the JTT matrix-based model [35] The tree with the highest log likelihood (-61359.70) is shown. Initial tree(s) for the heuristic search were obtained automatically by applying Neighbor-Join and BioNJ algorithms to a matrix of pairwise distances estimated using a JTT model, and then selecting the topology with superior log likelihood value. The tree is drawn to scale, with branch lengths measured in the number of substitutions per site. The analysis involved 17 amino acid sequences. All positions containing gaps and missing data were eliminated. There were a total of 3713 positions in the final dataset. Evolutionary analyses were conducted in MEGA7. A timetree was calibrated, specifing Um as the outgroup. To convert relative to absolute divergence time a divergencetime from 400-600 Mill was specified for the An to Um switch.

The timetree shown was generated using the RelTime method [36]. Divergence times for all branching points in the topology were calculated using the Maximum Likelihood method based on the JTT matrix-based model [1]. The estimated log likelihood value of the topology shown is −61359.70. The tree is drawn to scale, with branch lengths measured in the relative number of substitutions per site. The analysis involved 17 amino acid sequences. All positions containing gaps and missing data were eliminated. There were a total of 3713 positions in the final dataset. Evolutionary analyses were conducted in MEGA7.

### Gene homolog & dNdS analysis

Orthologous genes to all single copy genes were identified for all three proteomes by blastp comparisons (e-value: 10^−3^) against the single-copy families from all 12 species available from the FunyBASE. A two way BLAST between proteomes was performed to get the percentage identity and alignment parameters.

The orthologous gene pairs were globally aligned using t_coffee with default parameters and amino acids were replaced by codon from the cds sequence using the software pal2nal (<u>http://www.bork.embl.de/pal2nal/</u>) with options -nogap -nomismatch -output paml to generate a PAML input file. PAML was then used to calculate dN/dS in pairwise mode using the Yang and Nielsen method (Yn00 command). Protein sequences of *R. collo-cygni* were blasted e-value cutoff of 10^−10^ against protein database of *Z. tritici* proteins and vice versa. Only those genes with matches in both species were kept.

### SNP calling and summary statistics

Resequence data for all samples was mapped to our reference genome using SNP calling was performed using GATKs HaplotypeCaller [37] with ploidy set to 1. SNPs and indels were subsequently filtered for read and mapping quality. Vcftools [38] was used to calculate transition/transversion ratio and to convert the vcfs to ped files for further analysis. Data for SFS were directly extracted from the vcf files and visualised using ggplot [46]. Tajima’s D and nucleotide diversity were calculated on the genome and per gene using the package PopGenome [39].

### Population structure and phylogeny

To assess the population structure, we used the LEA package for R [40]. We analysed basic population structure by drawing a PCA for all data using the pca() function. We calculated the minimal cross entropy (mce) for the dataset. Mce is lowest when using low k values, thus we choose k = 2-4 for further analysis with the snmf() function. We performed 10 runs with alpha set to 10 and 100. Both analyses were run for both all SNPs as well as SNPs in coding regions only. Differences between all of these runs were minimal, thus only results for a single run (with alpha = 10, SNPs in coding regions only) were plotted.

To construct a population phylogeny we extracted all 162045 SNPs as alternative reference genomes using GATK and drew a Neighborhood Joining tree using PhyML in Seaview (GTR, NNI, BioNJ, 1000 bootstaps) [41,42].

### LD analysis

We calculated the recombination rate in ρ/kb using the softare LD helmet [43]. As LDHelmet requires long scaffolds to calculate breakage points, we performed the analysis on scaffolds 1-12. The software was run using the guidelines provided by the authors. We used a window size (-w) of 50, and recommended parameters for ρ searchspace. We specified a burn in of 100000 iterations and 1000000 true iterations. Block penalty (-b) was set to 50. The results for the upper and lower 2,5th percentile as well as the median were extracted from the binary results file and imported in R for visualisation. The histogram with ρ values per bin was plotted using ggplot2. LD graphs were plotted using the base plot function. These analyses were done once for all samples and once for a subset of the samples excluding the three outliers as identified by the phylogeny (AR, NO and NZ).

### Historic population size estimation

For the estimation of the historic population size (Ne) we used the software PopSizeABC [44]. We performed 200.000 simulations with each 50 independent fragments of 2Mb. Parameters for ρ were derived from the LDHelmet results, with a minimum cut-off at 1*10×^−8^ and a maximum at 1*10×^−7^ set as priors. Minimum allele frequency was set at 2 and the maximum Ne at 1000000. Summary statistics were extracted using the same parameters, with the tolerance, as recommended, set to 0.0005. *Ramularia* has a long latency period in the field and symptoms can only be observed near the end of the growing season, therefore we used generation time of 1 year.

## Results

### *Ramularia collo-cygni* is an emerging pathogen

To confirm the recent emergence of *R. collo-cygni,* we obtained seed samples from the Bavarian State Research Center for Agriculture (LfL) from all over Bavaria over the years 1958-2010. We quantified *R. collo-cygni* DNA from these archive samples dating back to 1958. This revealed that *R. collo-cygni* was constantly present in harvested barley grain (on average 39.4 pg R. collo-cygni DNA / ng total-DNA) (Figure 1). However, it is only starting 1984 that we observe irregular oscillations of enhanced DNA content in spring barley seeds (up to 710 pg R. collo-cygni DNA / ng total-DNA in 1984). In winter barley we observed even higher contents of *R. collo-cygni* DNA from 1989 on with maximum amount above 2000 pg *R. collo-cygni* DNA per ng total-DNA. These dates coincide with more frequent reports about epidemics and relative importance of RLS.

**Figure 1:**
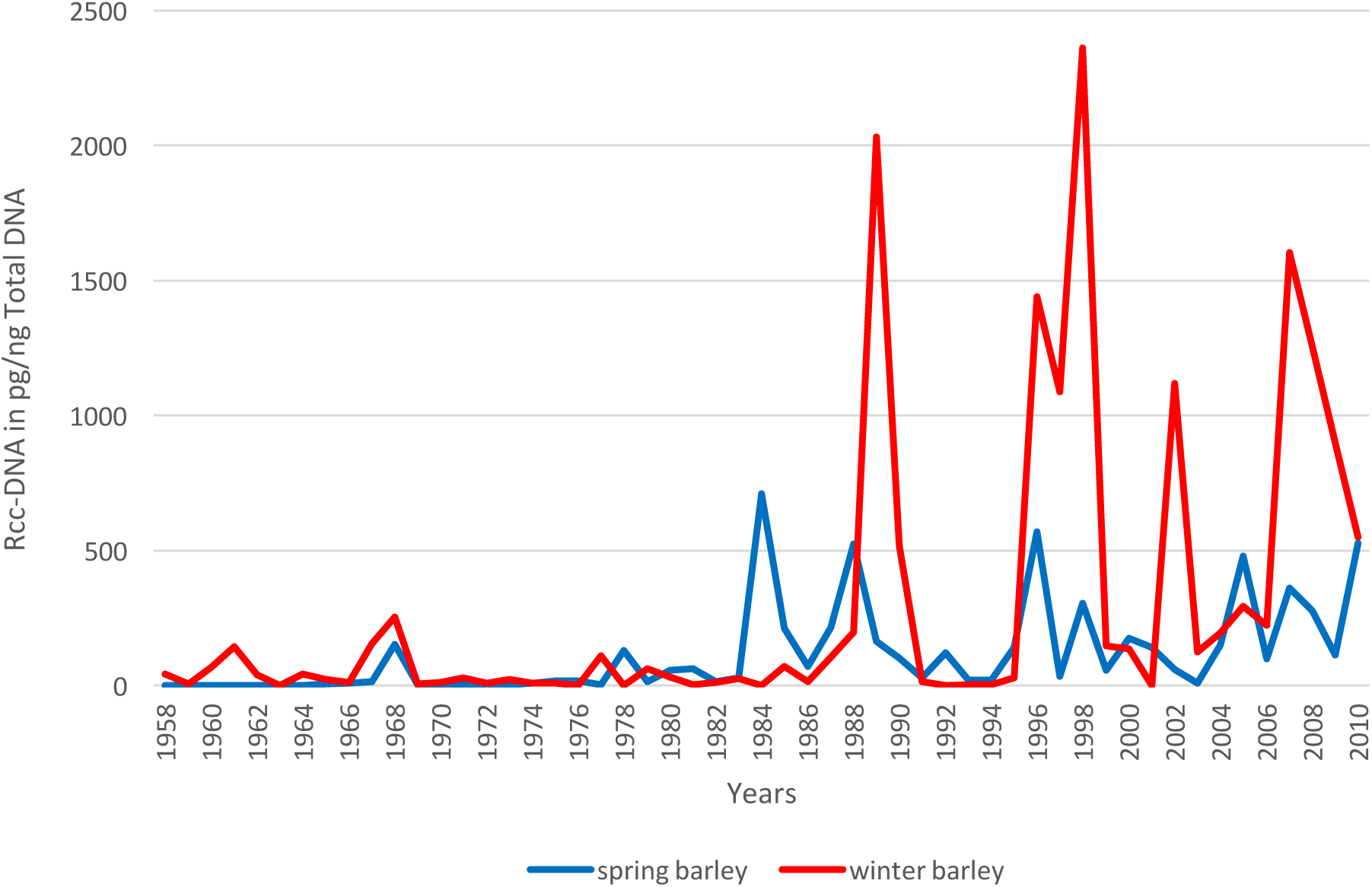
*Ramularia collo-cygni* DNA content in archive barley seed samples. DNA from barley seeds dating back from 1958 until 2010 was extracted. *R. collo-cygni* DNA amounts were calculated via quantitative polymerase chain reaction and are indicated in pictograms of *R. collo-cygni* DNA per nanograms of total DNA.

### On genome level *R. collo-cygni* is a typical Dothideomycete

We sequenced genomic DNA from *R. collo-cygni* isolate urug2 using a combination of a shotgun library on the Illumina MiSeq v3 and a long jumping distance library (LJD) on the Illumina HiSeq 2500 v3. This sequencing strategy allowed for sequence assembly into long scaffolds.

The assembled genome data of *R. collo-cygni* isolate urug2 contains about 32 Mb, in only 78 scaffolds with an N50 scaffold size 2.1 Mb (Table 1). Automated gene annotations are prone to misprediction of splice sites and actual coding sequences (CDS). Therefore, we performed RNA sequencing of mRNA isolated from *R. collo-cygni* isolate urug2 grown under six different axenic growth conditions to generate diverse sets of transcripts. Having these transcripts enabled us to re-annotate problematic splice sites and coding regions (CDS) of all genes, which we considered crucial for follow-up analyses. Using the RNA-seq data mapped to the genome, the annotation was manually corrected gene by gene yielding a more reliable set of gene models (Figure S1) shows an example of improved gene models). The curated *R. collo-cygni* annotation enabled the prediction of 11,637 protein coding genes. Out of these 11,637 predicted genes, 11,614 were expressed in at least one of the growth condition (Table S1). The use of RNAseq enabled us therefore, to achieve the most accurate annotation and a strong confidence in our gene models for 99.8% of them.

**Table 1.** Comparative genome statistics.

**1Table 1 Comparative genome statistics**
|  | *Ramularia collo-cygni* | *Ramularia collo-cygni* | *Phaeosphaeria nodorum* | *Pyrenophora tritici-repentis* | *Zymoseptoria tritici* |
| --- | --- | --- | --- | --- | --- |
| Isolate | ***Urug2*** | *DK05 Rcc001* | *SN15* | *Pt-1C-BFP* | *IPO323* |
| Genome size (Mb) | **32.3** | 30.3 | 37.2 | 38.0 | 39.7 |
| Chromosomes | ***n.d.*** | *n.d.* | *n.d.* | *n.d.* | 21 |
| Scaffolds | **78** | 576 | 107 | 48 | 21 |
| N50 scaffold (Mb) | **2.1** | 0.210 | 0.179 | 0.116 | *Finished* |
| GC-content (%) | **49.7** | 51.4 | 50.2 | 50.0 | 51.7 |
| Protein coding genes | **12,346** | 11,617 | 14,885 | 12,169 | 10,933 |
| Gene density<br>(Number of genes per Mb) | **383** | 384 | 402 | 329 | 276 |
| Total secreted Protein | **1,170** | 1,053 | 1,540 | 1,282 | 1,056 |

To get a better perspective of *R. collo-cygni* amongst and other fungi, We reconstructed a phylogenetic tree of *R. collo-cygni*, 12 closely and three more distally related species (Figure S2). We confirm previous findings that that *R. collo-cygni* falls within the *Mycosphaerellaceae* clade of the dothideomycete class [7,45] which contains important pathogens of different crops [54]. Additionally, we calculated the divergence time of the different pathogens in our phylogeny. These calculations show that *R. collo-cygni* (Rc) diverged from the closest sequenced relative (*Zymoseptoria tritici,* Zt) of 59 milion years ago, this is twice as long as the divergence time calculated between *Fusarium graminearum* (Fg) and *Fusarium fujikuroi* (Ff).

Interestingly, when general genome features are compared to other dothideomycete plant pathogens with available genome sequences (*Phaeosphaeria nodorum* SN15 [46,47], *Pyrenophora tritici-repentis* Pt-1C-BFP [48] and *Zymoseptoria tritici* IPO323 [49], *R. collo-cygni* doesn’t display any obvious unique characteristics in terms of genome size or composition (Table 1), although it shows many biological and phenological differences to these pathogens.

### *R. collo-cygni* coding and non coding sequences compared to other fungi

The fungal secretome is usually of pivotal nature in interactions with plant hosts [59]. The *Ramularia collo-cygni* secretome represents around 9% of its predicted proteome. We predicted 1292 secreted proteins ranging in length from 41 to 3256 amino-acids and 213 effector candidates genes (Table S2). This slightly exceeds previous annotation[13], but is still similar to the number predicted for related plant pathogens.

Next we compared *R. collo-cyngi* genes with 11 other dothideomycetes to look for unique and shared characteristics. A full genome comparison shows a clear bimodal distribution, indicating a large (>3500) conserved core-set of genes, shared between 10 or more species. Similarly, a second large fraction (>2800) genes is unique or only has a homolog in one of the other species. When counting the number of secreted proteins shared among these plant pathogens and endophytes (Table S3), very few (43) are shared among all the 12 species used for the analysis. 217 secreted proteins are shared between *R. collo-cygni* and one other species and 237 secreted proteins are unique (Figure 2A). The distribution of shared and unique genes changes even more when looking at the number of shared effectors candidates (Figure 2C). 43 effectors appear to be unique for *R. collo-cygni* and very few are shared among all species.

**Figure 2:**
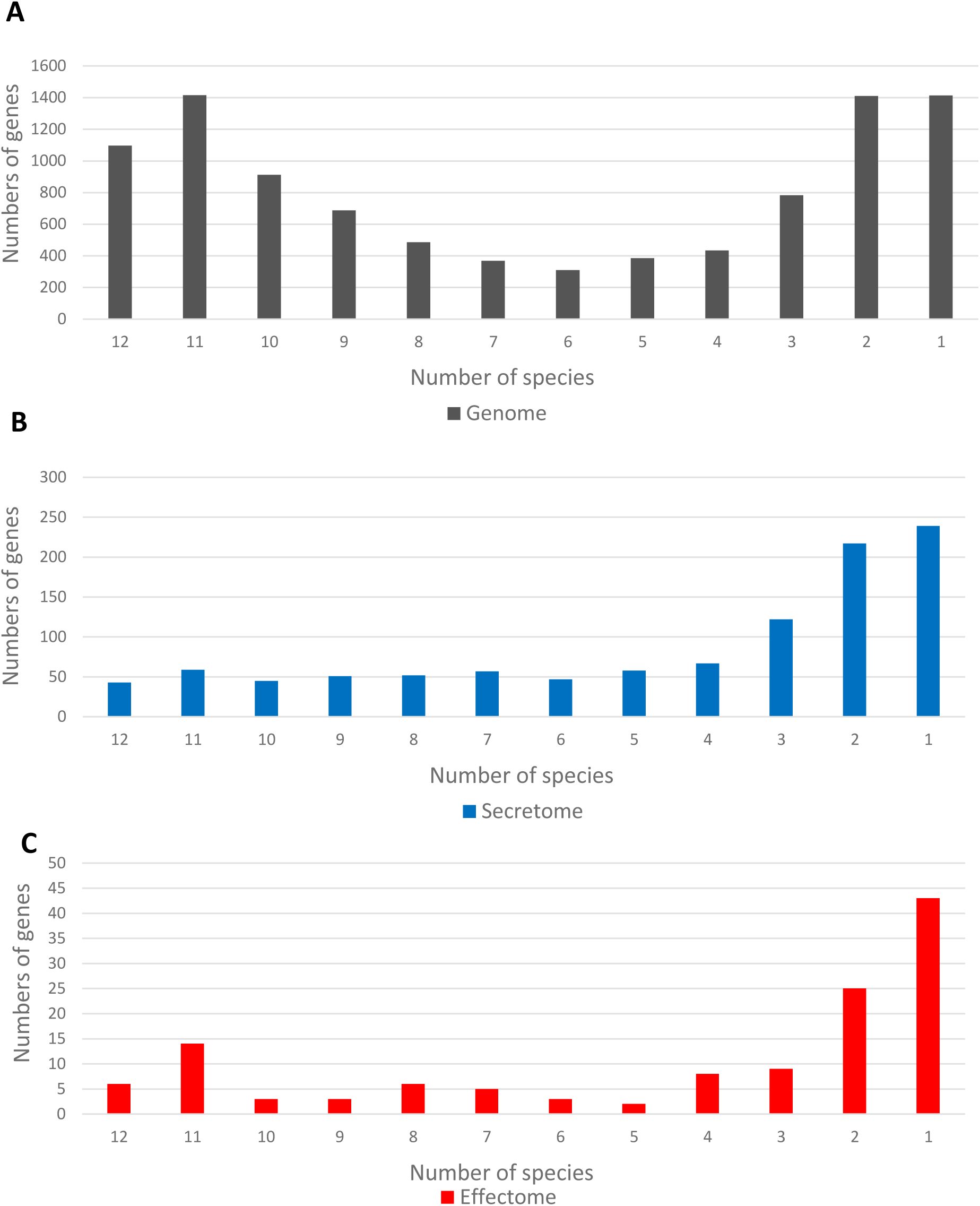
Numbers of homolog genes between *Ramularia collo-cygni* and 11 phytopathogenic fungi. Data is presented as histograms representing the numbers of homolog genes between *R. collo-cygni* and 11 or 10, 9, 8, 7, 6, 5, 4, 3, 2, 1, 0 other phytopathogenic fungi (*Fusarium graminearum; Pseudocercospora fijiensis; Magnaporthe grisea; Piriformospora indica; Pyrenophora tritici-repentis; Pyrenophora teres f. teres; Phaeosphaeria nodorum; Ustilago maydis; Zymoseptoria tritici; Blumeria graminis; Aspergillus nidulans).* **A**: Representation of the homolog genes in the whole genome. **B**: Representation of the homolog genes in the secretome. **C**: Representation of the homolog genes in the effectome.

Additionally, we compared the content of non-coding sequences and repeat sequences like DNA transposons and other transposable elements (Table S4). In terms of repeat sequences, *R. collo-cygni* ranks amongst the low end of the spectrum. Only 9.75% of the genome consists of repeats, whereas in *Rynchosporium commune* or the more closely related *Zymoseptoria tritici* are 31 and 21%, respectively.

### Strong divergence from *Zymoseptoria tritici*

To gain insight of the differentiation of *R. collo-cygni* from its closest related *Mycosphaerellaceae* species with high quality genome data, the wheat pathogen *Zymoseptoria tritici,* we calculated the ratio of non-synonymous substitution over synonymous substitution (dN/dS) (Figure 3, Table S5). The data shows high dS (dS>2), thus confirming that these two species have diverged a long time ago. Genes that are significantly differential expressed on Barley medium compared to other growth media show no higher signatures for selection (Figure 3A). When comparing putative secreted proteins and putative effectors shared between the two species, we find that the effectors show slightly higher dN/dS ratio (anova, p = 2.8*10^−8^) and a slightly different distribution as well (Figure 3B). Yet, in terms of absolute values and outliers, there are no clear outstanding gene candidates showing high dN/dS (Figure S3).

**Figure 3:**
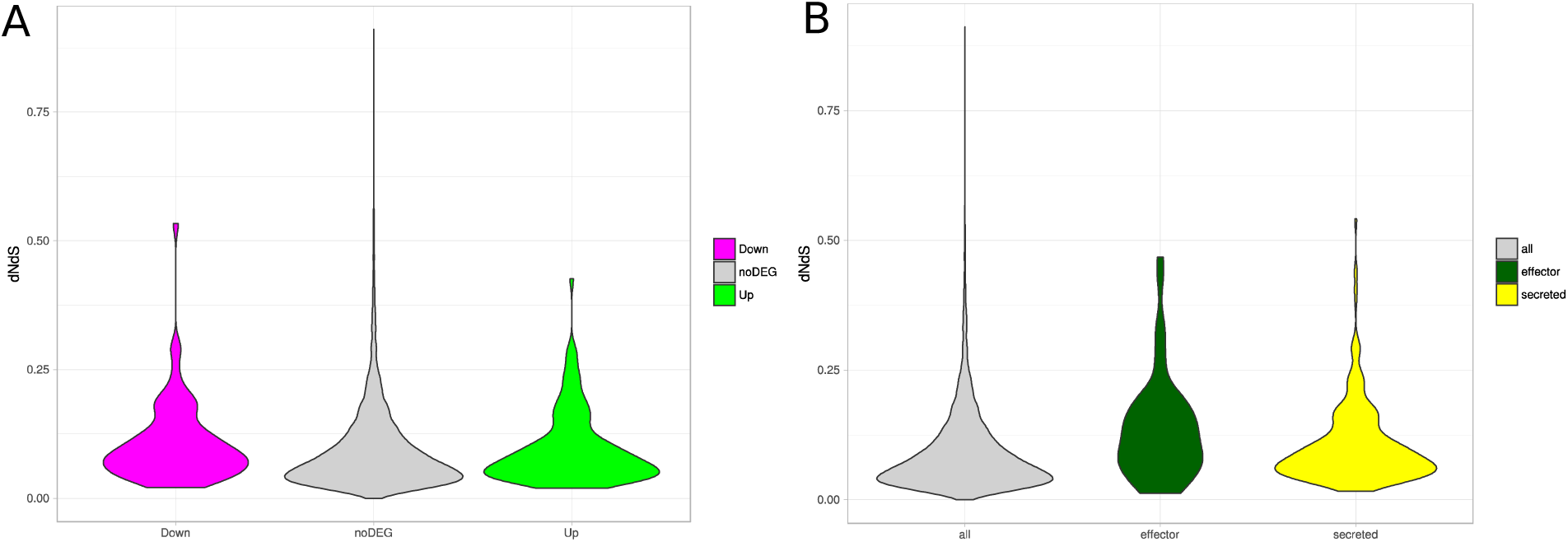
dN/dS between *R. collo-cygni* and *Z. tritici*. Violin diagrams of the dN/dS ratio for predicted proteins in a pairwise comparison between *R. collo-cygni* and *Z. tritici.*A) Data coloured according to respectively 2 fold Up regulation of Down regulation of the coding gene on BSA compared to the other growth media. B) Data coloured based on whether the proteins are predicted to be putatively secreted proteins or effectors.

### Sequencing of further *R. collo-cygni* strains

Based on the fact that *R. collo-cygni* diverged from other Dothideomycetes such as *Z. tritici* millions of years ago, but did not acquire truly outstanding genome characteristics, or genes under selection in our genome comparisons, we set out to investigate the population structure within the *R. collo-cygni* species in order to unravel its recent evolutionary history. We used whole genome sequencing data for 14 isolates from different geographical locations and five from different host plants. All reads from the sequenced isolates were mapped back to our reference genome and SNPs were called.

In total we obtained 162045 high confidence SNPs between all samples of which 79949 SNPs occurred within coding regions, corresponding to approximately one SNP every 200 bases. The transition/transversion ratio is 2.36 for the whole genome and 2.31 for the coding sites, suggesting no anomalies in the SNP calling. 56% of all SNPs in the coding regions occur as singletons within the population (Figure S4). Average nucleotide diversity per site (π) over all coding sites is 4.7 × 10^−4^ and Tajima’s D is −1.186. When looking at per gene statistics, median value of π per site is 4.7 × 10^−4^ and Tajima’s D is −1.1630. These values as well as there overall distributions remain very similar when the data is split in expressed and non expressed genes or in the secreted protein and predicted effector fractions (Figure S5). There are no significant differences between any of the groups. Such overall low diversity, high singleton rate and low Tajima’S D is indicative of recent population expansion.

### *R. collo-cygni* isolates show little substructure in the global population

Several approaches were used to assess the population structure of *R. collo-cygni*. First, we reduced the complexity of the data and drew a PCA for the SNPs. Figure S6 shows that the first two components explain 17% (12 and 5 resp) of the variation. When drawing the two first components, most isolates group very closely together in one cluster. Only three outliers can be observed. Similar patterns are observed in the coding parts of the genome only as well in the whole genome. Next we estimated the population structure from the SNP data. The previous PCA results and minimum entropy analysis suggest the optimal cluster k-value to be between 1 and 3 (Figure S7). LEA[40] was run for k-values 2-4 in 10 separate runs. Figure 4A shows that with these k-values, two or three isolates can be marked as outliers, being the isolates from Norway, Argentina and possibly New Zealand. All other samples group together with k = 3 and show a transient pattern for k = 4. No segregation of isolates can be seen, neither on a geographical basis nor based on the host of origin.

**Figure 4:**
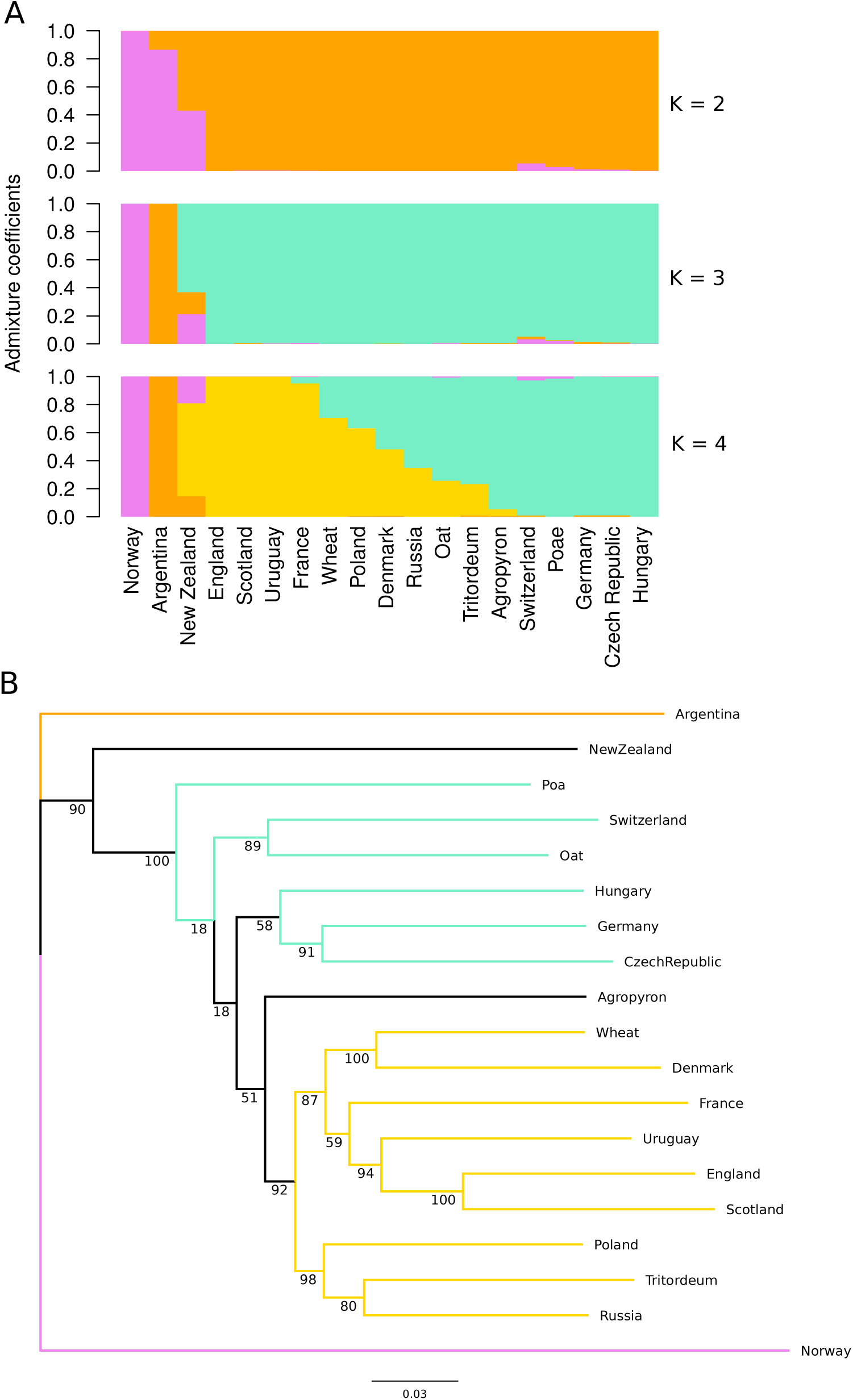
R. collo-cygni shows limited subtructure. A) Admixture plots for k + 2-4 show limited population structure. Three outlier samples can be defined, one of which is an admixture between all three “groups”, yet all other samples show different degrees of admixture without resolving either host or geographical properties. B) Phylogenetic reconstruction the the 19 resequenced samples shows similar population (non) structure. Two clear outliers can be observed (AR, NO). A large number of the internal branches show low bootstrap values, suggesting limited support which corresponds to likely admixture of the samples. Branch colors represent the colors of part A.

To confirm these findings, we extracted all high confidence polymorphic sites from reconstructed genome sequence for each of the isolates. The resulting alignment was used to construct a phylogeny using PhyML (NNI, 1000 bootstraps). Figure 4B shows that indeed isolates from Norway and Argentina are the two main outliers, followed by New Zealand. Both figures possess the same color coding and show indeed very similar internal structures. The low bootstrap values at some of the internal branches confirm that no clear subgroups can be defined within the inner branches of the tree. Thus, except for three outliers, there is no clear population structure for *R. collo-cygni* in our dataset.

### Signatures of recombination indicate sexual reproduction

Next we wondered whether there are signatures of sexual recombination within the species as suggested following marker analysis. We used LDHelmet [43] to estimate the linkage disequilibrium within the species on the longest scaffolds of our assembly (> 250 Mb). Figure 5 shows that when analysing all 19 isolates, rare, but very clear LD events can be observed. ρ values across the genome are mostly lower than 10^−6^/kbp, however, in small windows these values change up to 8 (Figure S6). When we repeated the LDHelmet analysis without the three aforementioned outliers, these peaks in ρ almost disappear (Figure 5). Indicating that in fact the majority of our samples might originate from an asexual population.

**Figure 5:**
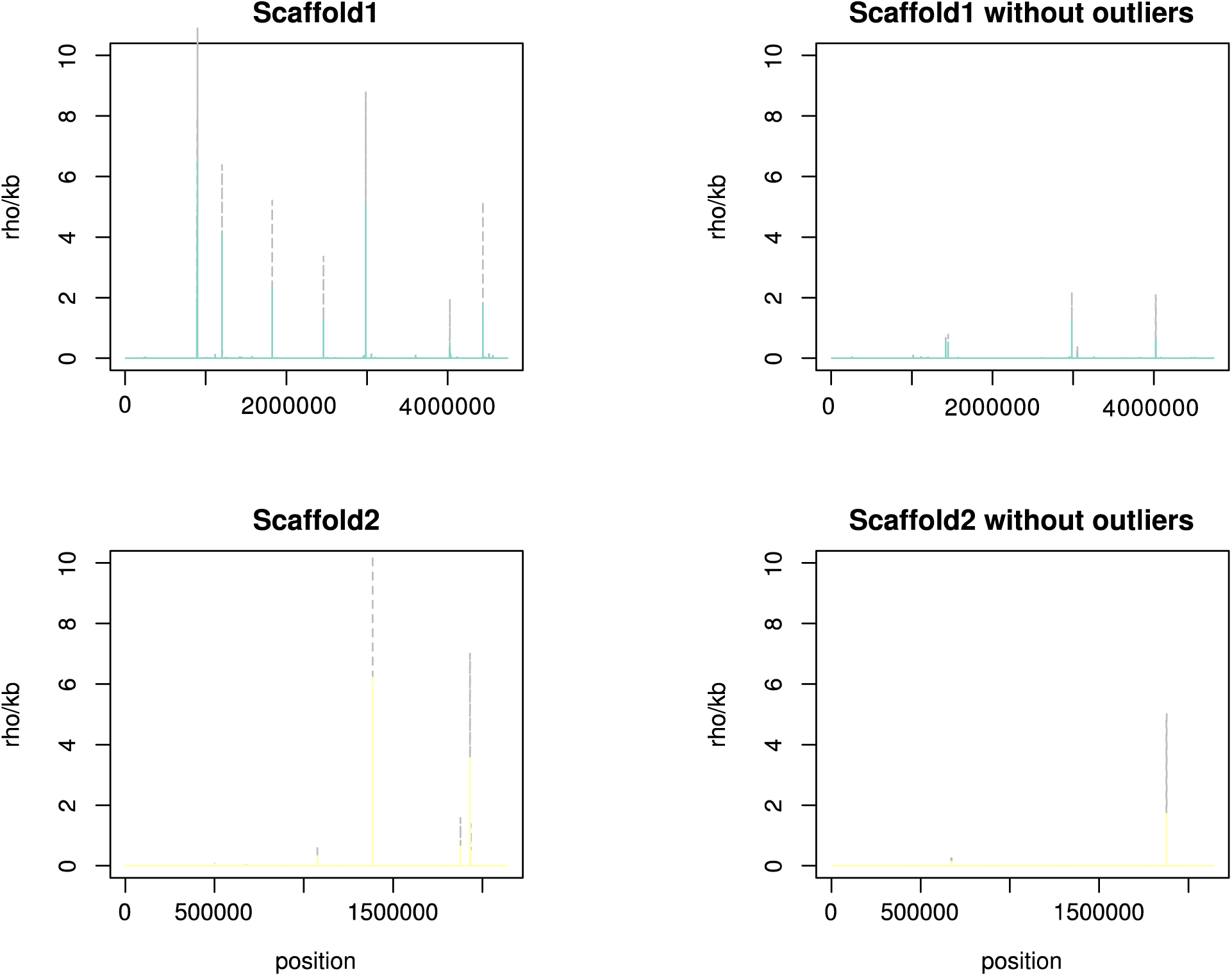
Recombination can be observed in R. collo-cygni. A) Recombination in rho/kbs (y-axis) was calculated per position on the scaffold for the 12 longest scaffolds (>1.000.000 bp). Scaffold 1 and 2 are shown. Other scaffolds can be found in S. Figure 7 Left shows the result for the for all samples, right the results excluding the three outlier samples. The x axis shows the scaffold length in bps

### Recent expansion of the effective population size

To understand the current *R. collo-cygni* epidemics we wanted to gain better insight in the current and historical effective population size (Ne). We can take advantage of the high quality genome assembly and the high length of the genomic scaffolds and use the Bayesian Computation package PopsizeABC [44] to calculate Ne back in time. We calculated the required summary statistics using data from the first 12 scaffolds and performed 200.000 simulations to estimate the historic Ne p to 13000 generations ago. This shows that *R. collo-cygni* has undergone a bottleneck between 1000 and 150 years ago after that the Ne increased 100-fold to about 100.000 (Figure 6).

**Figure 6:**
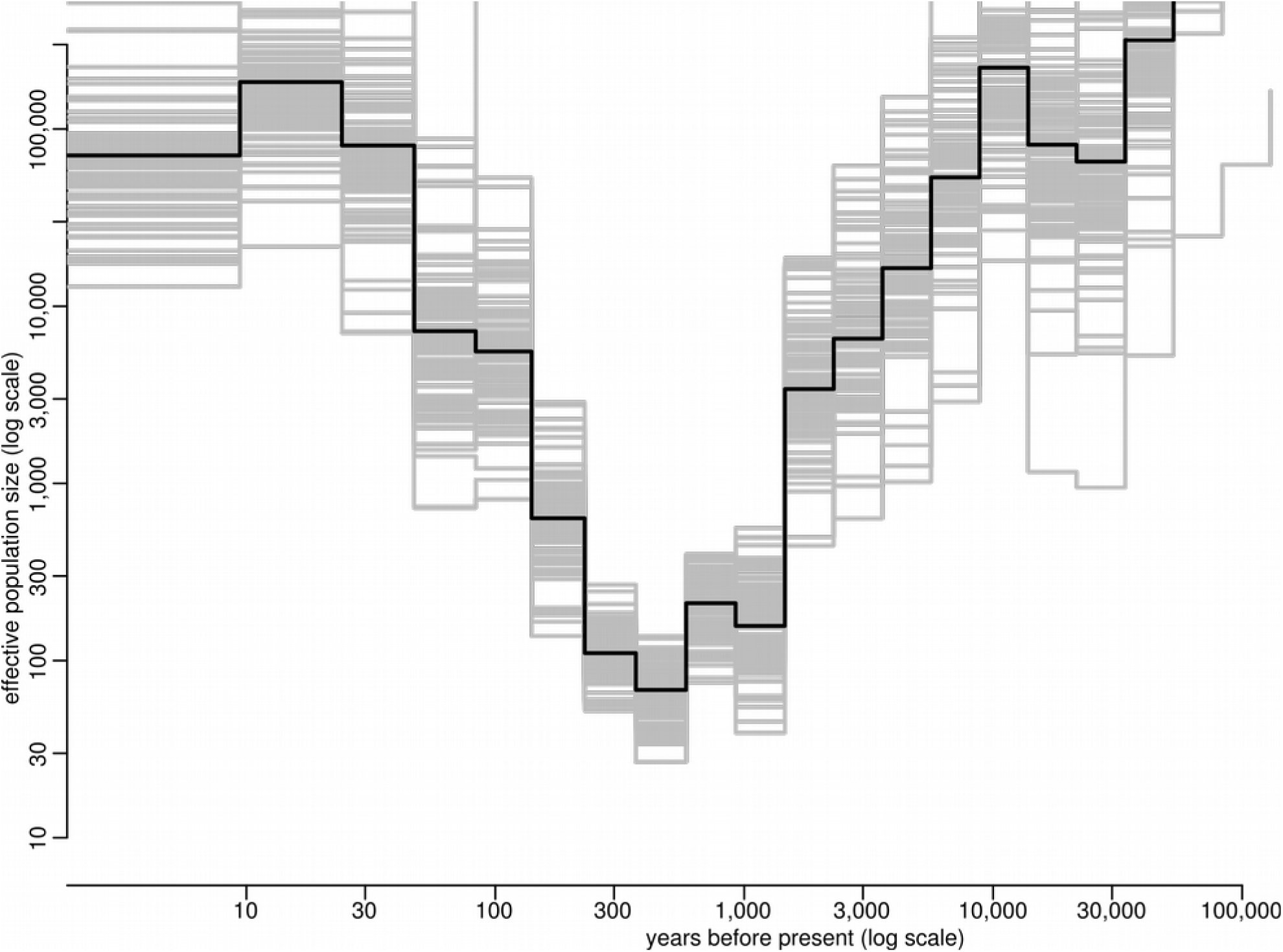
R. collo-cygni experience gradual Ne expansion. Overview of estimated historic population size back in time. Expected Ne was estimated by comparing observed summary statistics with 200.000 simulations using PopSizeABC. Minor variations could be observed for each run. Grey lines represent 100 individual runs and the plack line shows the median values for those.y-axis shows the effective population size (Ne) agains the number of years (or generations) ago (x-axis).

## Discussion

Although, it was discovered more than 100 years ago as a saprophyte of grasses [6], *Ramularia collo-cygni* began to attract interest only since the late 1980’s when it started to gain an economic impact on cultivation in regions of major barley production. Our analysis of barley seed archive samples dating back to the late 1950’s showed that the fungus was already present in grain before epidemics were recognized. However, it is only since the mid-1980’s that there is a significant increase *of R. collo-cygni* DNA in the grain. Indeed, coinciding with more frequent observations of RLS symptoms from the field [1].

A draft genome of *R. collo-cygni* (isolate DK05) has been available since 2016. The data suggested a genetic composition that might at least partially explain the lifestyle of *R. collo-cygni*, which is characterized by a long endophytic phase throughout the life cycle of the host and an intense parasitic phase during crop senescence [7]. We generated an independent draft genome for another isolate (urug2) to get better insights in *R. collo-cygni* variability and understand the genomic factors that contribute to its current success as a pathogen. To allow robust population genetics analysis, we aimed for optimised gene annotations and increased scaffold length. We assembled the 32 Mb genome into only 78 scaffolds of N50 2.1 MB length with 12,346 high confidence genes predicted. Overall, our sequencing approach allowed us to produce a high quality draft assembly for comparative studies and confirmed that unlike many other pathogens [50] *R. collo-cygni* did not undergo any genome expansions since it diverged from it’s nearest related sequenced *Dothidiomycete* species 59 million years ago.

One main feature of pathogenic fungi to avoid or suppress plant defenses is the secretion of the so-called effectors. Unlike what can be seen between certain oomycete species, where the numbers of genes in some effector families differ in order of magnitude within the genus [51] *R. collo-cygni* effector numbers are comparable with related fungi. Moreover, this phenomenon is often associated with high repeat content on the genome [52], yet *R. collo-cygni’s* repeat content is low. Hence such gene family expansions are unlikely to explain the differences between these species.

When looking for similarity between the predicted *R. collo-cygni* proteome, secretome and effectome to those fungal genomes see a very strong bimodal distribution with large numbers of genes shared between all species, indicating highly conserved genes, but we find also small sets of (effector) genes unique for each pathogen. As there is no reported race-specific full resistance in *R. collo-cygni*, selection on the shared effector genes might not be expected to be the dominant driving force for pathogen evolution.

To test that hypothesis we performed a pairwise comparison of the coding sequences of *R. collo-cygni* and the related wheat pathogen *Z. tritici*. We observe a very high synonymous mutation rate and see very little evidence for strong positive selection on certain gene-types between the two species. The dN/dS ratio has a simple and intuitive interpretation of selection pressure, but comes with a large number of limitations, especially when dS is high [53]. For this reason, our findings are not conclusive with regards to the number of genes actually under selective pressure. However, our analyses provide interesting insights. Contrary to phenomenon observed in a large number of other plant pathogens, we see no evidence for accelerated evolution of secreted proteins or effectors. The dN/dS distribution is similar for all three classes and also when looking at pathogenicity-related genes, based on up or downregulation on BSA, a host, mimicking medium, there are no significant differences between the groups. This is in stark contrast to for example the genus *Colletotrichum,* where high dN/dS was visible in effectors when comparing endophytic and parasitic species [54] or the comparison between two *Phytophthora* species, where an elevated (>1) dN/dS was much larger in the effectors than in all genes together (37% versus 14%)[55] and positive selection on effectors appear to drive host-adaptation [56].

This leaves the possibility that only a few unique effectors are shaping the evolution of *R. collo-cygni.* In other fungal pathogens, such unique effectors are often associated with host specificity or aggressiveness on a certain host, as for example in *Z. tritici* [52] or *B. graminis* [57,58]. Moreover, appearance or disappearance of effector genes might indeed contribute to co-evolution with the plant hosts and is a potential driver for speciation. Thus the previously mentioned effectors unique to *R. collo-cygni* might contain interesting candidates for future analysis. Alternatively, the largely symptomless and endophytic lifestyle and the possibility to escape to alternative hosts might have caused little selection pressure on *R. collo-cygni*.

Next, to obtain a better understanding of the biology and the current world-wide emergence of *R. collo-cygni*, we analysed 19 *R. collo-cygni* strains. This revealed a relatively low SNP rate, as well as a very similar transition/transversion ratio on coding regions and the whole genome (2.31 and 2.37 resp.). These findings are the first indication of gradual, but not extreme diversification within the species and suggest a bottleneck and recent expansion event of the species. We confirm that indeed there is very little evidence for geographical population substructure and with the exception of three possible outlier samples, all our samples form one genetic cluster. In a previous study, populations from Czechia and Scotland were found to form two diverged populations [16]. With our additional sampling, we now show that isolates from these populations form rather two sides from within the same large main cluster but not clear outliers. The main cluster can be divided in two putative genomic groups, but these do not show a strong geographical cline. These results are partly surprising, because other cereal pathogens, like for example *Wheat Yellow Rust fungus Puccinia striiformis f.sp. tritici,* show geographical population structures [59].

One explanation for lack of clustering and mixing could be long distance dispersal of spores. Wind dispersal could be one means by which *R. collo-cygni* spores travel around the globe [60,61], such dispersal is heavily reliant on stochastic processes [62], but that would only in part explain the lack of geographical clustering, as geographically close areas are still most likely to be genetically more similar to one another. Another reason could be found in seed transmission and contaminated seed shipments by humans. Seed transmission of *R. collo-cygni* has been suggested before [63]. Hence, trading and winter nurseries potentially contribute to dispersal of *R. collo-cygni* whereas seed monitoring and quarantine might delay global spreading of RLS. These finding are in line with observations made for another barley pathogen with seed dispersal, *Rynchosporium commune* [64], but closely related pathogens like Z*. tritici* are rarely isolated from seed [65].

Interestingly, our analyses also show no sign of a clear host specialization. These results are in contrast with other fungi. The rice blast fungus *Magnaporthe oryzae* is pervasive on rice, but strains can also be found on many other monocot hosts [66], *Z. tritici* contains a small set of genes that can be related to host preference [52] and even the broad host range, opportunistic pathogen *Botrytis cinerea* shows signs of host-adaptation in its population structure [67]. These examples are all linked to either selection of or absence/presence of effector genes. Yet, since we see no clear evidence for such phenomena in *R. collo-cygni,* it will be crucial to identify other factors in *R. collo-cygni’*s biology that determine its opportunistic properties. This is particularly relevant, because other wild graminaceous hosts can be an alternative source of inoculum, resulting in more than one overwintering host available that provides a “green bridge” or “seed bridge” for *R. collo-cygni* to survive winter.

The lack of host adaptation on the fungal side could give further evidence for *R. collo-cygni* being a general endophyte of grasses. Pathogenicity would then derive from host specific lack of resistance factors or presence of susceptibility factors, likely triggered or at least enhanced by environmental factors. In *Piriformospora indica* metabolic cues from its hosts result in differentstrategies of host-colonisation by directly affecting expression levels of genes related to specific life-styles [68]. Similar specific interactions are described for symbiosis [69].In fact early infection with *R. collo-cygni* has been observed to have beneficial effects on yield, suggesting that the host-parasite interaction switches from mutualistic/endophytic to parasitic [70]. An early symbiontic interaction could also explain the lack of major resistance genes in barley. The absence of epidemics in related *Gramineae* and the high diversity in barley germplasm give hope that effective host genes can be identified for the control of RLS.

Rapid expansion of a plant pathogen can also arise from beneficial recombinations during sexual reproduction. Using LD scans, we show that indeed there is proof of sexual recombinations between the populations we inspected. These findings confirm previous assumptions [15,16]. Interestingy, there is limited recombination within the main subpopulations (or main isolate cluster), that excludes Argentinia, Norway and New Zealand, showing that the recent expansion of *R. collo-cygni* is possibly due to clonal propagation of one or several well-adapted strains. The combination of high clonal reproduction and global transfer via seed was probably favoured by modern intensification of cereal production and global cereal trade and trafficking. When calculating the effective population size, we see a strong increase in the past 200 years, coinciding with the intensification of barley production, suggesting that *R. collo-cygni* expanded over time with the increased availability of susceptible hosts.

With its large efective population size (10 times larger than that of *Z. tritici* [71]), mixed recombination and global gene flow *R. collo-cygni* meets the criteria to classified a high risk pathogen [72]. To date no major R genes have been identified against *R. collo-cygni* in barley and the characteristics described above make that eventual R genes are unlikely to provide durable resistance, even when introduced as stacks or pyramids [73]. The best way to diminish the current epidemic is to drastically reduce the effective population size. This could be achieved by measurements to reduce overwintering capacity and seed transfer or more mixed breeding approaches including identification of host factors that result in quantitative resistances or manipulation of host susceptibility factors.

## Acknowledgements

This work was financially supported by grants from the Bavarian State Ministry of Food, Agriculture and Forestry (Projects KL/12/07 – and KL/08/07) and the Bavarian State Ministry of the Environment and Consumer Protection in frame of the project network BayKlimaFit (to RH, subproject 10,). We thank C. Wurmser (Chair of Animal Breeding TUM) for support with sequencing. Special thanks goes to C. Hutter and R. Dittebrand for technical assistance. Without their patience and accuracy isolating, maintaining and extracting the fungal cultures the study would not have been possible.

---

**Figure S1.**
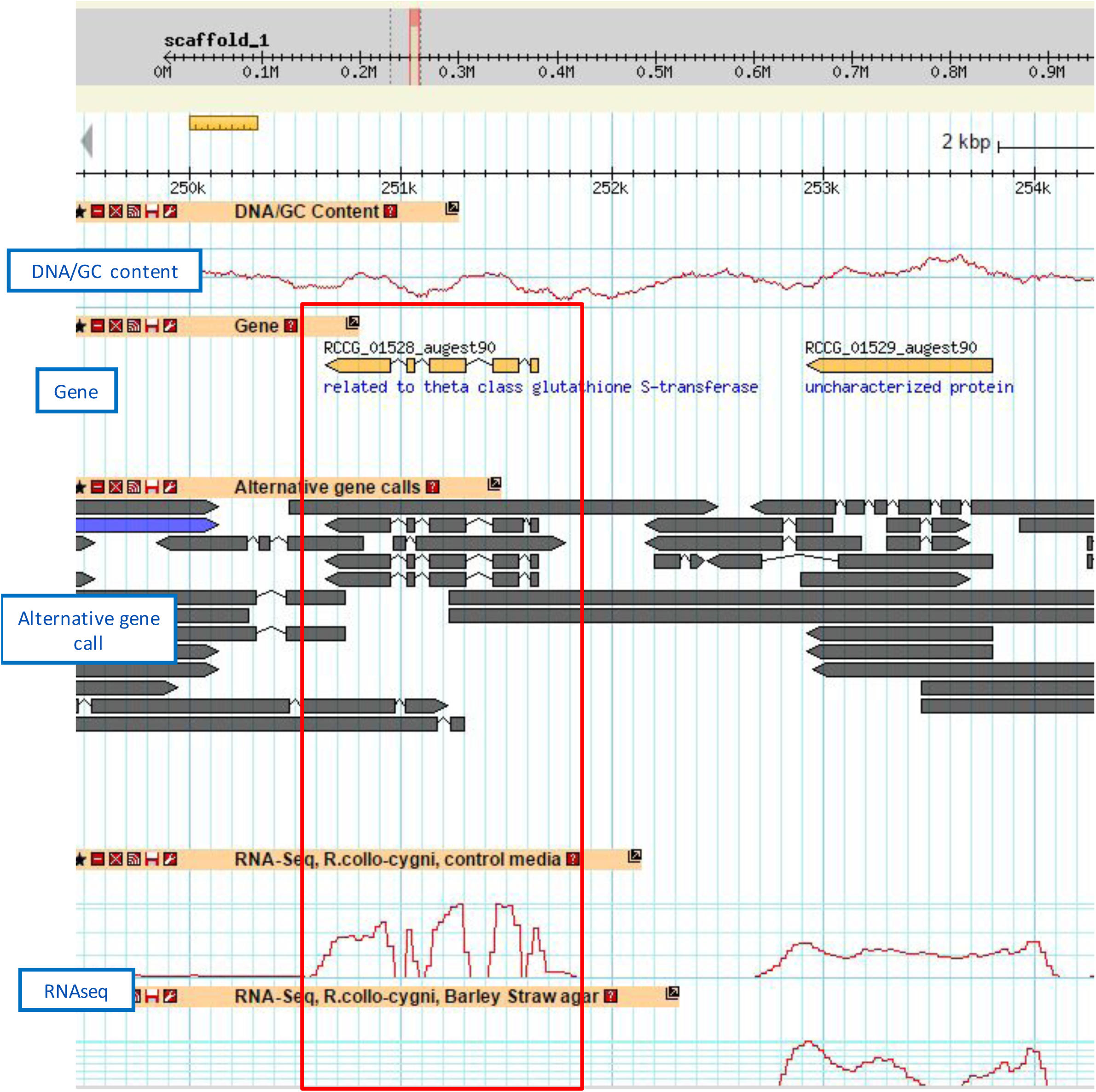
Genome browser screenshot showing the alternative gene model called using gene predictors and the accuracy that the RNAseq brings in choosing the right model.

**Figure S2.**
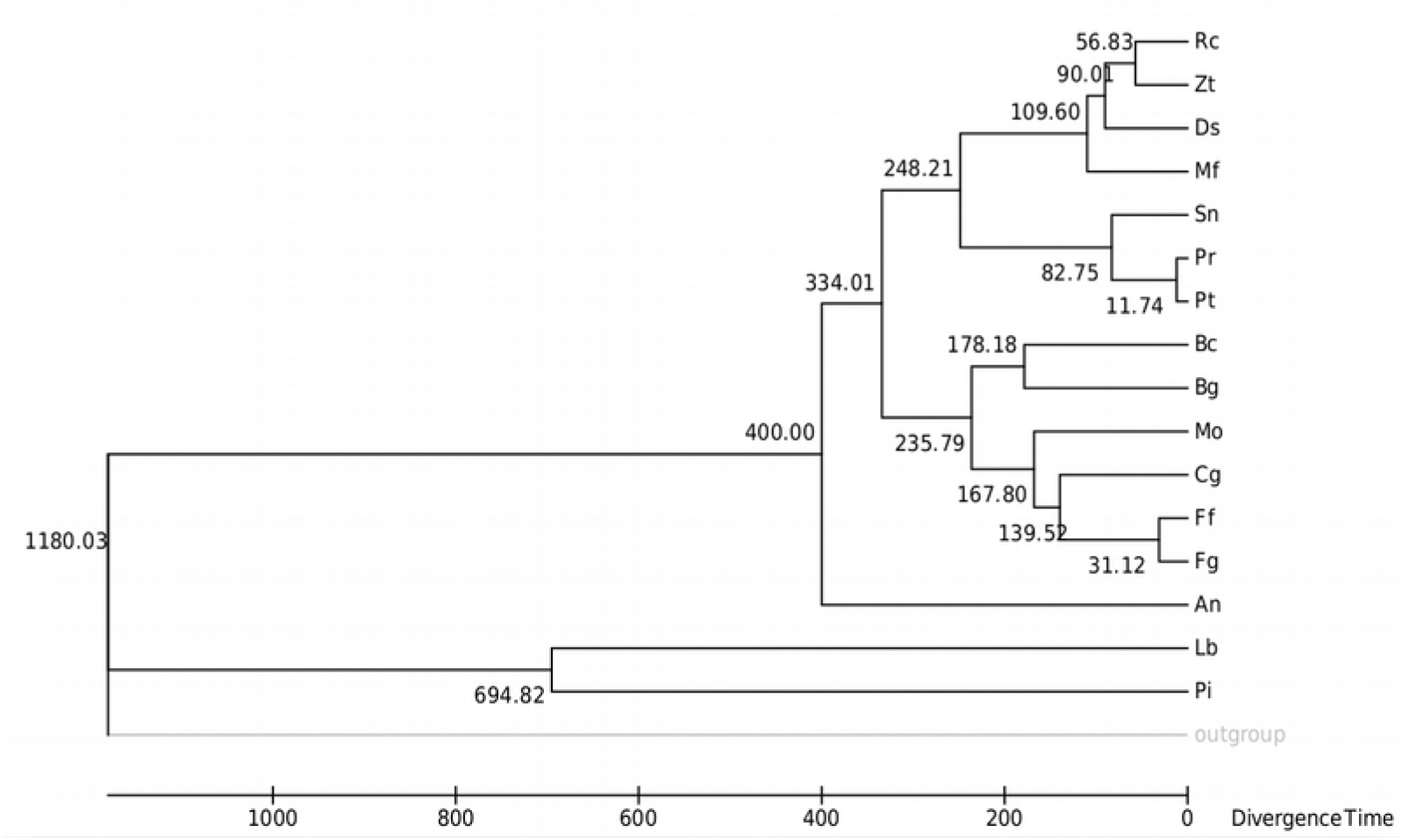
Phylogenetic tree showing the relationship between Ramularia collo-cygni and 16 other fungi. Data is presented as a maximum likelihood phylogenetic tree based on the analysis of 10 housekeeping genes. Cg = *Colletotrichum graminicola*; Ds= *Dothistroma septosporum;* Fg= *Fusarium graminearum;* Mf= *Pseudocercospora fijiensis;* Mo=*Magnaporthe grisea;* Pi= *Piriformospora indica;* Pr= *Pyrenophora tritici-repentis;* Pt= *Pyrenophora teres f. teres;* Rc= *Ramularia collo-cygni;* Sn= *Phaeosphaeria nodorum;* Zt= *Zymoseptoria tritici;* Bc= *Botrytis cinerea;* Bg = *Blumeria graminis;* An= *Aspergillus nidulans;* Lb*= Laccaria bicolor;* Ff= *Fusarium fujikuroi*.

**Figure S3.**
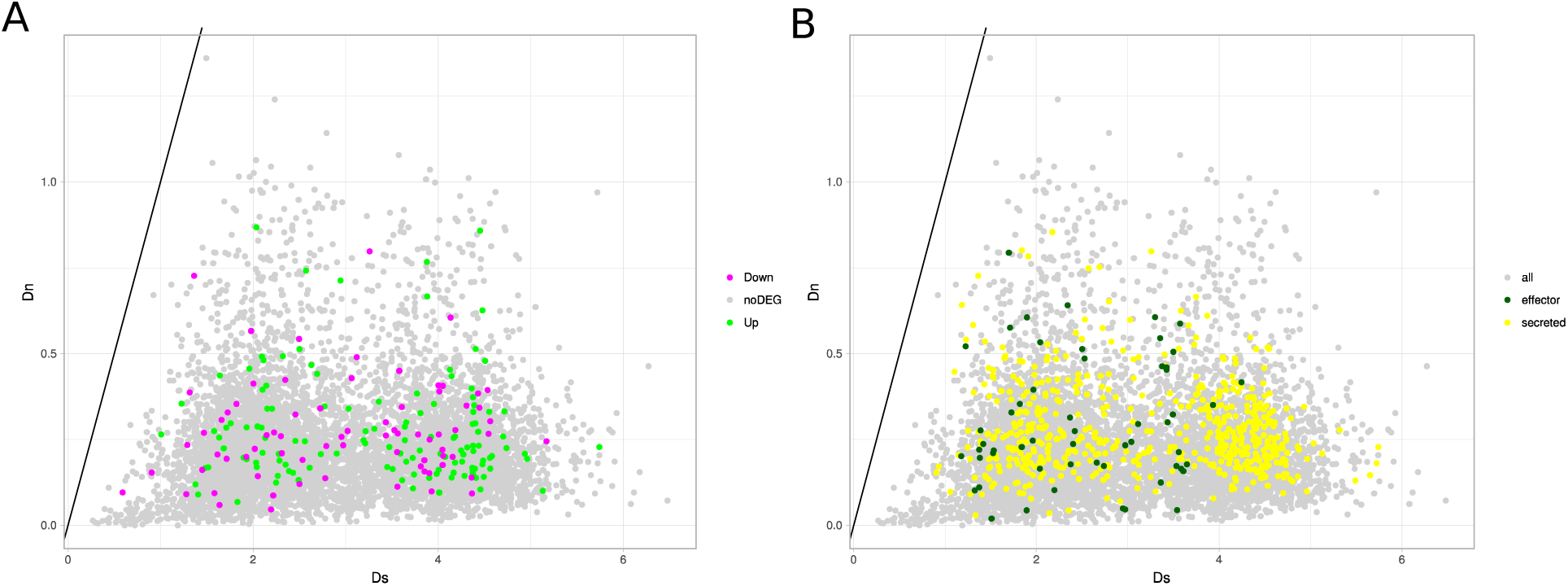
Scatter plots of the Ds (x axis) agains Dn (y axis) for each predicted protein in a pairwise comparison between *R. collo-cygni* and *Z. tritici.* A) Data coloured according to respectively 2 fold Up regulation of Down regulation of the coding gene on BSA compared to the other growth media. B) Data coloured based on whether the proteins are predicted to be putatively secreted proteins or effectors.

**Figure S4.**
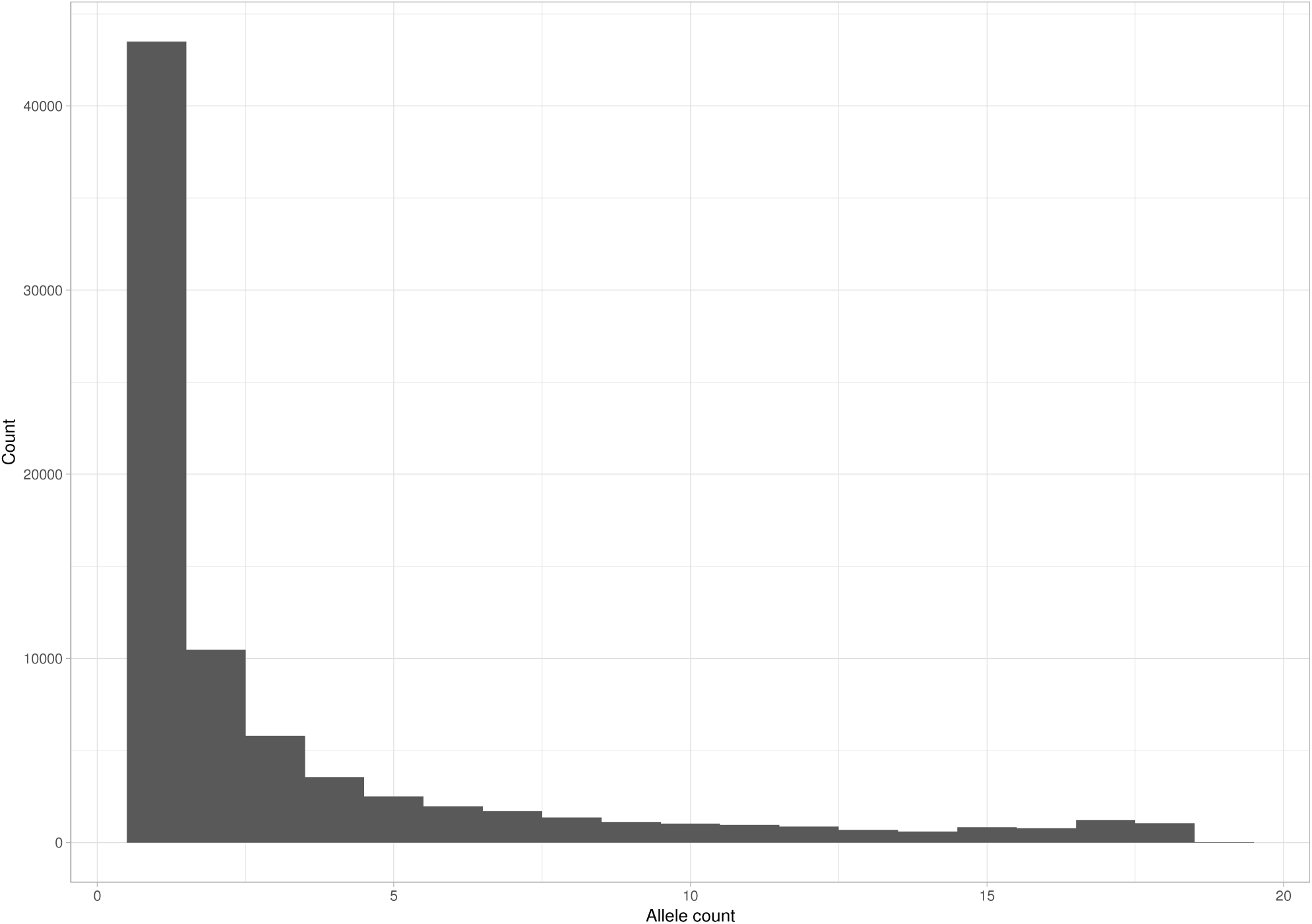
Site Frequency Spectrum for all SNPs in the coding regions between 19 *R. collo-cygni* isolates. The majority of SNPs are singletons, only very few SNPs show intermediate or high allele frequencies.

**Figure S5.**
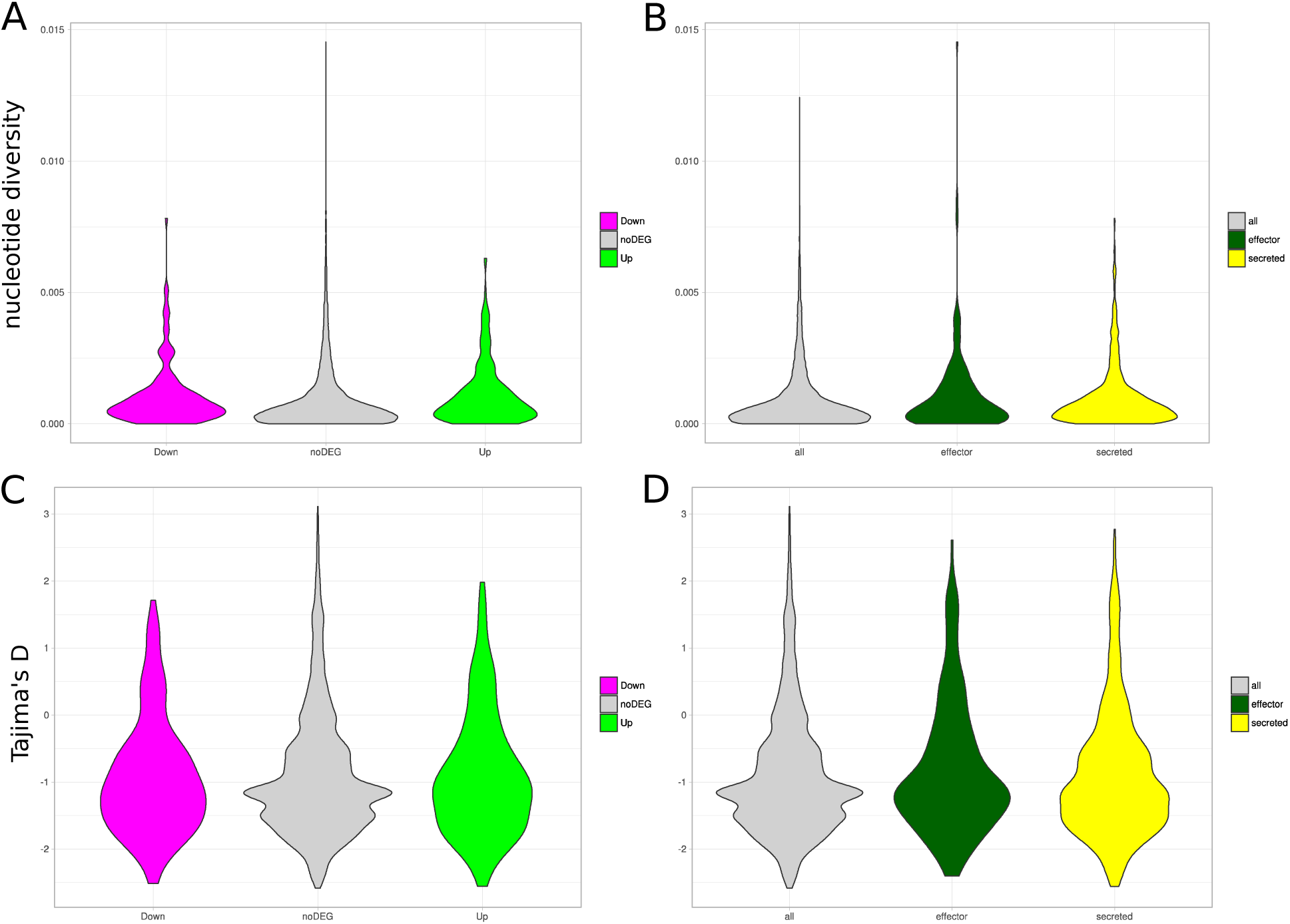
Violin diagrams of A,B) the nucleotidediversity and C,D) Tajima’S D per gene for all genes in the *R. collo-cygni* genome based on our world-wide collection. A,C) Data coloured according to respectively 2 fold Up regulation of Down regulation of the coding gene on BSA compared to the other growth media. B,D) Data coloured based on whether the proteins are predicted to be putatively secreted proteins or effectors.

**Figure S6.**
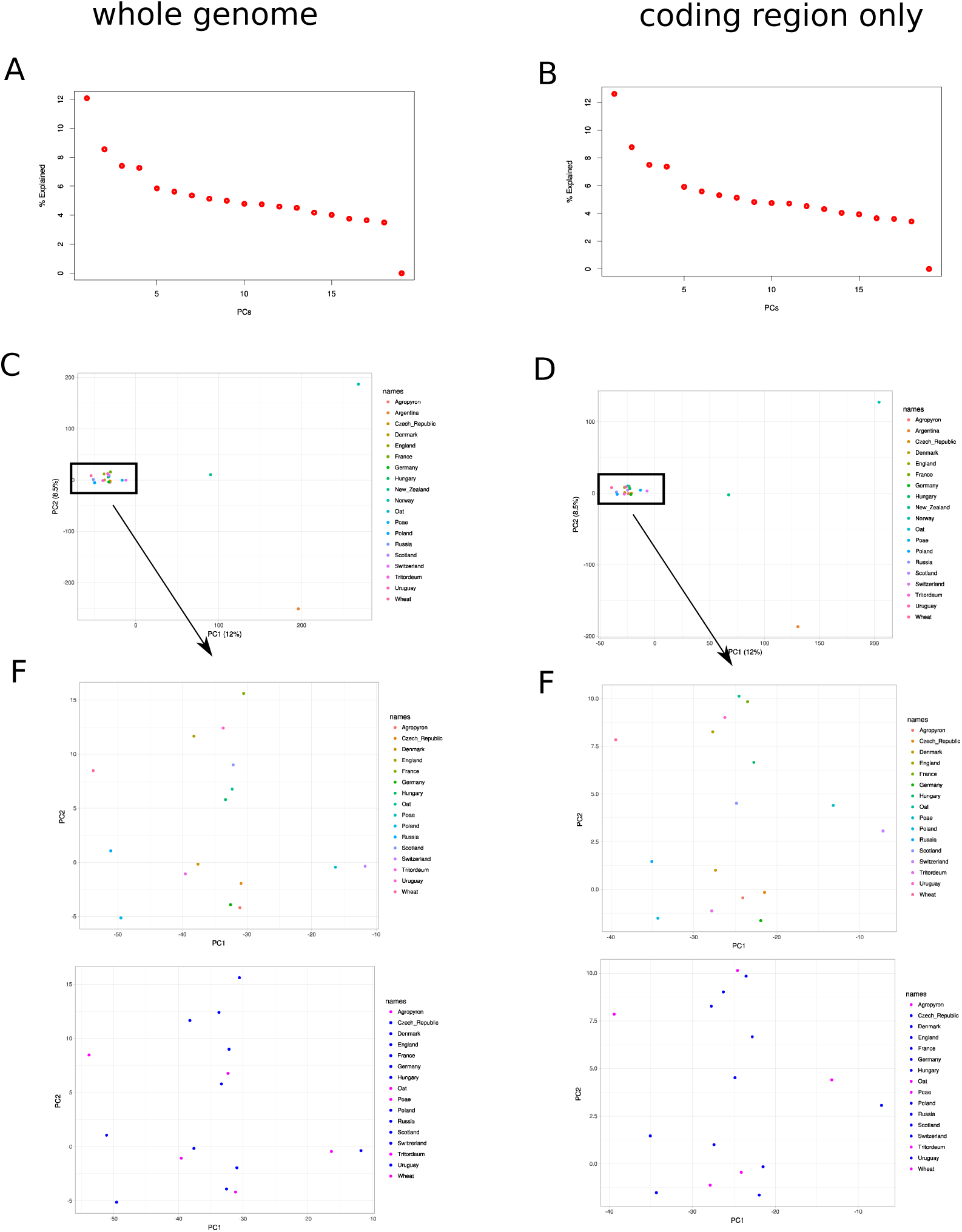
A,B) Eigenvalues of the calculated principal components (PC) for the whole genome and the coding regions only. The Y axis explains the % of variation that can be explained by each PC and the axis ranks the PCs from 1 to 19, with 1 and 2 having the most explanatory power. B,C) PCA for the first two principal components show one cluster for 16 samples and three outlier samples for the whole genome and the coding regions only. E and F show a close-up for the main cluster with the top panels colored by origin. All alternative host samples originate from Switzerland. The lower two panels colored by true host (barley) or alternative host. No substructures can be observed in either of the graphs.

**Figure S7.**
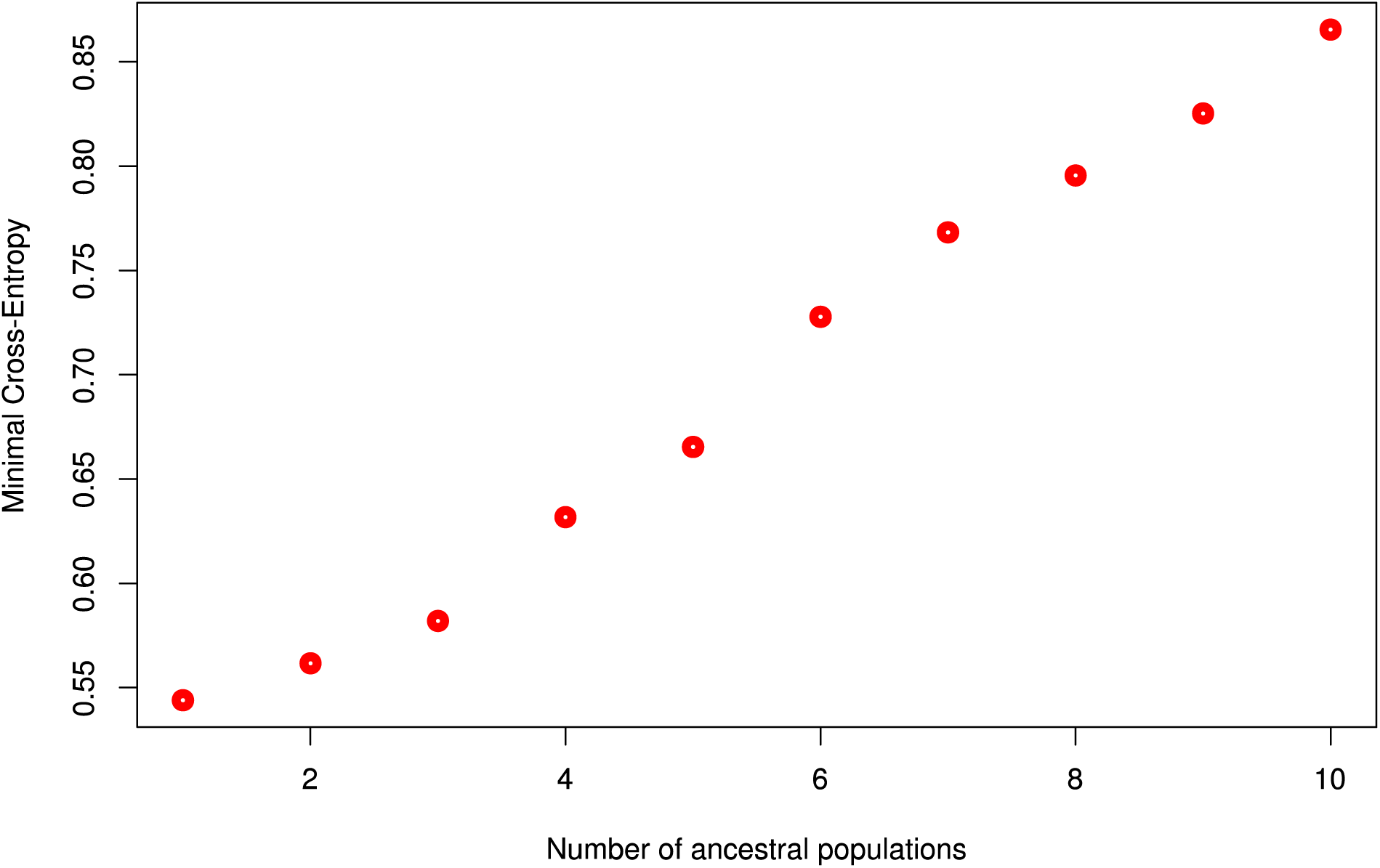
Minimum Cross-Entropy as tested for different number of ancestral populations (k-values) prior to running LEA. The larger the number of ancestral populations the larger the minimal cross-entropy.

**Figure S8.**
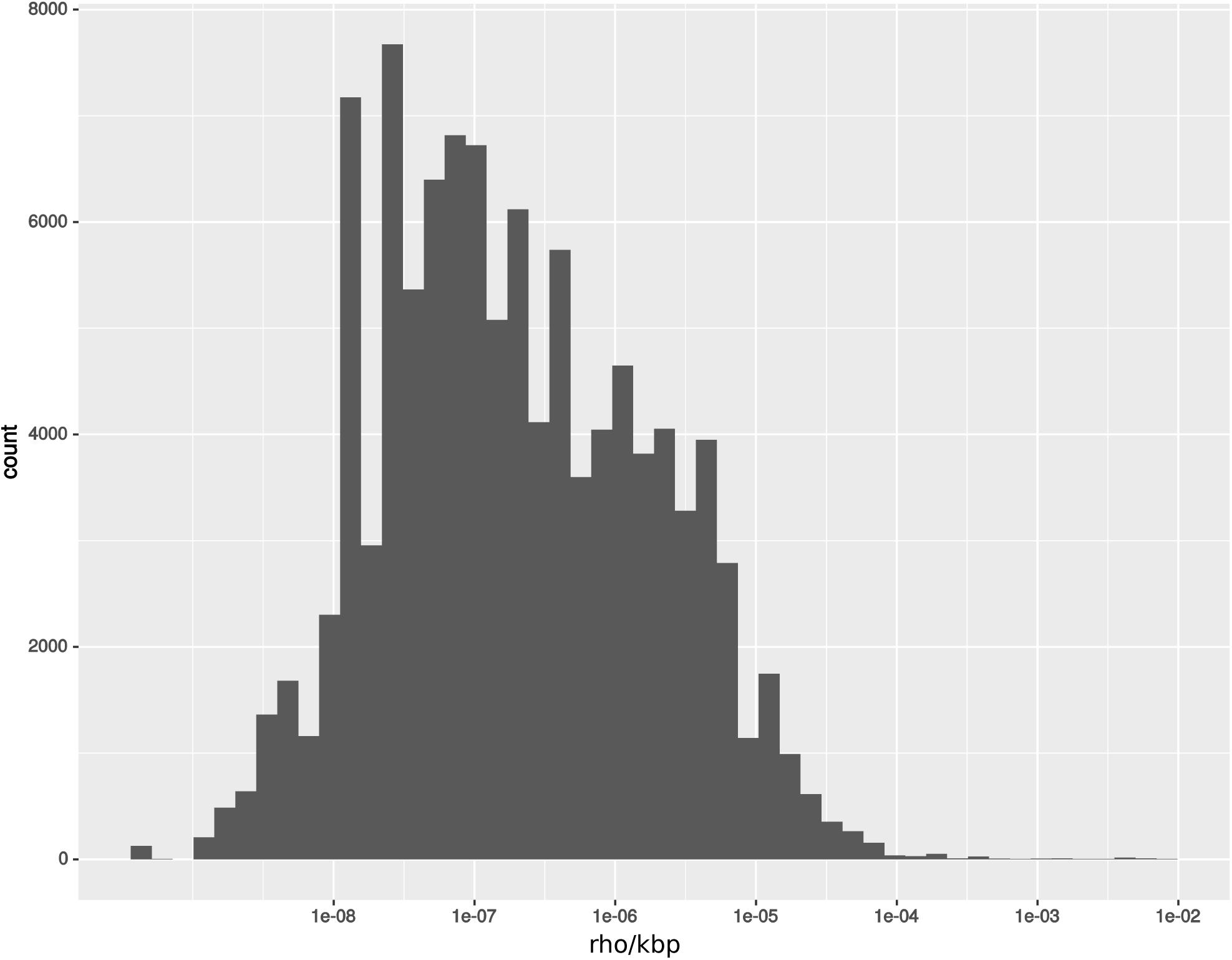
Histogram showing rho for each bin on the genome. X axis shows the different values and y axis the counts. Overal values range from 1×10^−8^ to 1×10^−5^, with outliers all the way up to 1×10^−2^.

**Figure S9.**
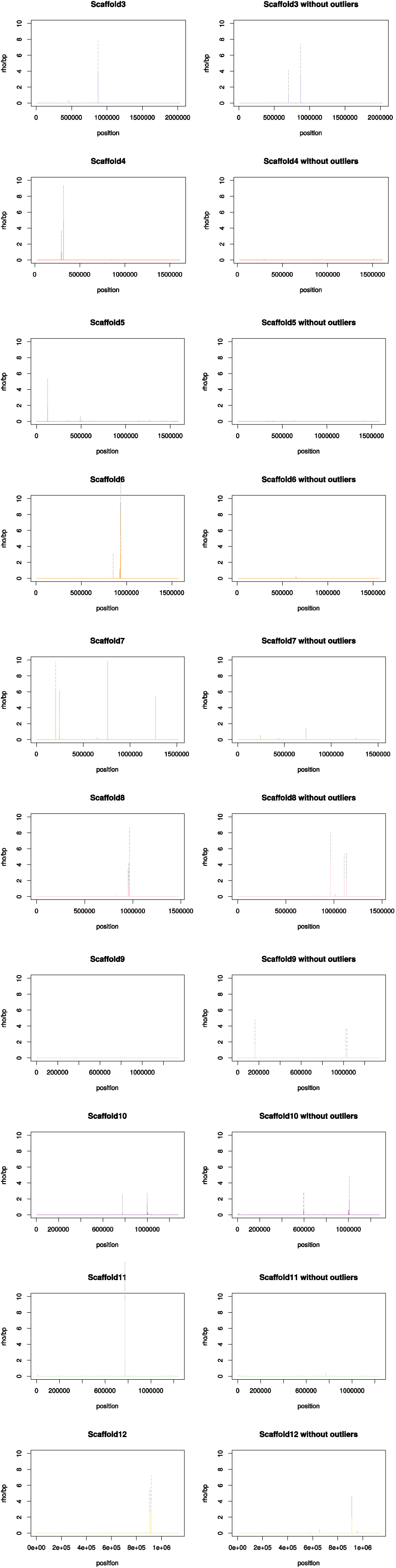
A) Recombination in rho/kbs (y-axis) was calculated per position on the scaffold for the 12 longest scaffolds (>1.000.000 bp). Scaffolds 3-12 are shown. Left shows the result for the for all samples, right the results excluding the three outlier samples. The x axis shows the scaffold length in bps

